# A comparative brain atlas in the Mexican cavefish identifies widespread changes in cellular composition and gene expression

**DOI:** 10.64898/2026.05.12.723532

**Authors:** Kathryn Gallman, Edward Ricemeyer, Aakriti Rastogi, X Maggs, Emilio Méndez Scolari, Erik R. Duboue, Nicolas Rohner, Harini Iyer, Wes Warren, Alex C. Keene

## Abstract

Understanding how naturally occurring genetic variation shapes human health and disease is critical for improving diagnosis and treatment strategies. The Mexican cavefish, *Astyanax mexicanus*, represents a powerful system for evolutionary medicine, enabling investigation of naturally evolved mechanisms of resilience to disease-related traits including diabetes, obesity, insomnia, and eye loss. Larval *A. mexicanus*, like zebrafish, are transparent, allowing whole-brain imaging, circuit mapping, and the generation of computationally derived atlases that precisely quantify neuroanatomical differences between surface and cave populations. Developing a molecular map of brain cell types provides a foundation for identifying evolved differences in neural circuits and physiology. Here, we present a single-cell atlas of the larval cavefish brain that reveals widespread divergence in the abundance and molecular signatures of neurons and glia. Our cell type map validates known neuroanatomical differences, including a reduction of the optic tectum and expansion of the pineal gland in cavefish. We uncover substantial changes in multiple glial cell classes that are linked to neural regulation of behavior, including microglia. Analysis of differential gene expression between surface and cavefish microglia revealed enhanced genes associated with synaptic pruning and clearance of neural debris, suggesting cavefish increased microglia activity to shape brain development. We also analyzed cell types that did not classify as canonical neurons or glia and identified notable divergence in transcriptomes and cell composition, including reduced meningeal fibroblasts in cavefish and substantial transcriptional changes related to phototransduction in non-visual photoreceptors within the pineal gland. Together, these findings provide a comprehensive atlas of cell type-specific gene expression differences between *A. mexicanus* surface and cavefish, establishing a platform for dissecting the molecular and cellular basis of evolved disease resilience in cavefish

## Introduction

A relatively small number of genetic models including *C. elegans*, *Drosophila*, zebrafish, and mice, have been central to uncovering fundamental genetic processes contributing to diverse aspects of brain function [1–3]. Large-scale genetic screens in these organisms have demonstrated that core molecular and neural processes governing behavior are deeply conserved across animals [4,5]. However, most classic genetic models rely on highly inbred or isogenic laboratory lines. This genetic homogeneity limits the ability to investigate how naturally occurring genetic variation shapes biological traits, trait interactions, and adaptive or resilient phenotypes [6]. In addition, phenotypes are often examined in isolation, despite many biological traits emerging from interactions among multiple physiological and neural systems. These limitations highlight the need for complementary work in outbred lines and genetically diverse animal models.

Extending functional interrogation beyond zebrafish through the use of a growing number of fish models provides additional avenues for studying foundational biological processes [7,8]. These systems include species that are uniquely adapted to extreme environments including the turquoise killifish which lives in ephemeral rainwater pools and ages through an entire lifespan in a single season, as well as African cichlids, which have reduced resource competition through rapid speciation and niche exploitation [9,10]. The genetic specializations in these particular systems provide a natural framework to answer questions related to aging and rapid evolutionary radiation. The availability of chromosome-level genomes and gene-editing technologies across many novel aquatic models has enabled the identification of genes and pathways underlying complex, evolutionarily derived traits, complementing and extending insights gained from traditional genetic model systems [11,12].

The Mexican tetra, *Astyanax mexicanus*, represents a particularly powerful emerging model for studying the genetic and evolutionary basis of human disease–relevant traits [13–15]. This species consists of an eyed surface population inhabiting rivers of Northeastern Mexico and southern Texas, and at least 34 blind cave populations that independently converged on similar, cave-adapted phenotypes. Many of these convergent traits are relevant to human disease phenotypes, including sleep loss, diabetes-like insulin resistance, obesity, fatty liver, reduced heart regeneration, eye degeneration, and autism-like behaviors [16,17]. *A. mexicanus* populations are easy to maintain, produce large broods, have transparent larvae suitable for functional imaging, and have high-quality genome assemblies and tools that enable diverse genetic manipulations [18,19]. Together, these features position *A. mexicanus* as a powerful system for advancing our understanding of the neural regulation of behavior and its relationship to disease.

Across species, ranging from fruit flies to mice, computationally derived anatomical brain atlases have been essential for defining the cell types that govern brain function [20–22]. These atlases rely on high-resolution imaging of brains labeled with cell type specific markers, followed by computational registration and averaging [22]. This approach has been especially successful in zebrafish at six days post-fertilization (6 dpf), when larvae are transparent and whole-brain imaging can be performed in intact animals [23–25]. These neuroanatomical atlases, when combined with markers of neural activity, have provided unprecedented insight into the relationship between brain structure and function in a vertebrate system. Building on these approaches, we have generated multiple comparative brain atlases of surface and cave populations of *A. mexicanus*, revealing widespread neuroanatomical differences and region-specific changes in neural activity [26,27]. However, a limited understanding of cellular diversity in the *A. mexicanus* brain remains a major barrier to interpreting these anatomical, functional, and behavioral differences.

Single-cell atlases have been used extensively to define neural cell types in aquatic organisms [28–31]. Zebrafish atlases, in particular, have provided high resolution mapping of cell types to function [24,31–33]. Beyond zebrafish, comparative atlases across related aquatic species have yielded insight into how cell type composition and transcriptional programs diverge during evolution. Although a single-cell atlas of the adult *A. mexicanus* hypothalamus revealed extensive differences in gene expression and cell type abundance between surface and cave populations, no whole-brain atlas currently exists for this species [27,34]. Developing a comparative whole-brain atlas at 6 dpf is critical to provide context for functional experiments, including neural ablation, activity imaging, and behavioral assays linked to the evolution of brain function.

Here, we present the first comparative single-cell atlas of *A. mexicanus* in an early-developing brain. Using single-nuclei RNA sequencing (snRNA-seq), we profiled dissected brains from 6 dpf surface fish and Pachón cavefish. We identified and characterized neuronal, glial, and additional non-neuronal cell populations across the brain. Our analyses revealed widespread differences in both cell type composition and gene expression profiles between surface and cave populations. This atlas uncovers evolved differences in brain cell types and provides a foundational resource for functional interrogation of genetic and cellular mechanisms underlying behavioral and disease-relevant traits.

## Methods

### Fish husbandry and maintenance

*A. mexicanus* were maintained under standard conditions and in accordance with protocol guidelines approved by the Texas A & M Institute for Animal Care and Use Committee [35]. Fish were housed at 23°C under a 14:10 light/dark cycle and fed tetra min flakes once daily. The surface fish were initially derived from Rio Choy and have been described previously [36]. The Pachón population was obtained from Dr. Bill Jeffery (Maryland) and has previously been described [36]. Breeding was induced in stock populations by increasing water temperature to 28°C while feeding a high-calorie diet of bloodworms multiple times per day [37]. Eggs were collected the following morning and raised under standard conditions in 250 mL of water in glass bowls, 12 cm x 6 cm (Pyrex 7200) with 50 larvae per bowl until 6 days post fertilization (dpf). *A. mexicanus* do not feed until 6 dpf, and therefore, larvae were not fed prior to dissections.

### Brain Dissections

Larval *A. mexicanus* aged 6 dpf from either the Río Choy surface population or the Pachón cave population were anaesthetized on ice immediately prior to dissection between Zeitgeber Time (ZT) 4 and ZT6. Brains were dissected in ice-cold water and placed into sterile 1.5 mL tubes on ice until flash-frozen in liquid nitrogen. Tubes were flash-frozen with approximately 100 larval brains per tube. This resulted in a total of four surface fish tubes and four cavefish tubes where each tube was treated as a single sample and used as a biological replicate for downstream statistical analysis.

### Nuclei isolation, library preparation, and sequencing

Flash-frozen samples were processed by Singulomics (SingulOmics Corporation, New York, USA) using the 10x Genomics platform (10x Genomics; Pleasanton, CA). Briefly, nuclei were isolated from thawed samples and used to construct 3’ single cell gene expression libraries using the 10x Genomic Chromium System (Next GEM v3.1). Libraries were then sequenced on an Illumina NovaSeq (Illumina; San Diego, CA) with approximately 400 million paired-end, 150 base pair, reads per sample. Sequencing data processing: The AstMex3_surface annotated reference genome from NCBI (accession GCF_023375975.1) was used for library alignment via CellRanger (10x Genomics; Pleasanton, CA) [38]. After computing raw counts matrices for each sample using default parameters, we generated filtered counts using cellbender remove-background v0.3.0, also with default parameters [39]. We then used the scanpy platform v1.9.8 to perform standard clustering analysis[40]. All per-sample count matrices were aggregated together for quality control filtering that removed cells with greater than 40,000 detected genes, greater than 20,000 counts, or greater than 2 percent of counts attributed to ribosomal genes (Supplementary Figure 1). Counts matrices were then normalized and log scaled to account for differences in sequencing depth between cells and stabilize expression variance. Next, we selected for highly variable genes and regressed out counts per cell driven by differing library sizes. After preprocessing, we performed principal component analysis (PCA) on the scaled count matrices and used the cell-level PCA embeddings to integrate across samples using Harmony (Fig S1A; Korsunsky et al., 2019). Next, we ran leiden clustering on the nearest neighbors graph, computed from the integrated data set with a resolution of 1.2, and visualized the results using UMAP (Figure 1A).

**Figure 1:**
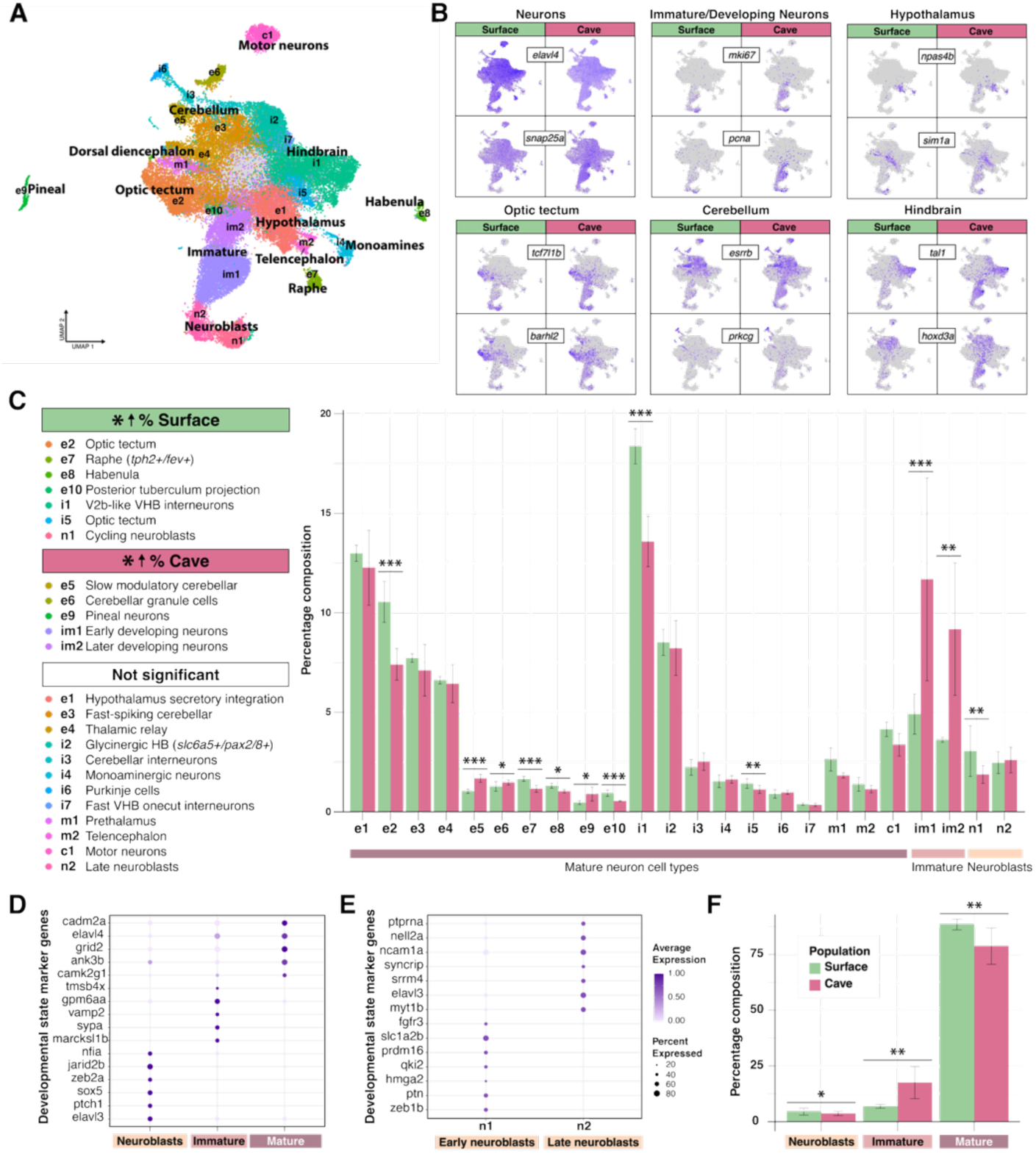
CNS neuron composition varies between *A. mexicanus* populations by neuron cell type and developmental stage. **A)** UMAP of 27 neuronal cell types labeled as excitatory (e), inhibitory (i), mixed excitatory and inhibitory (m), cholinergic (c), immature, (im), and neuroblasts (n). Label overlay provides context for developmental stage and regional location of mature neurons. **B)** Marker gene distribution for neuron identity, maturity, and regional locations show overlapping, but variable expression between surface and cavefish. **C)** Specific neuron cell types vary in percentage composition of the brain between *A. mexicanus* populations. Cell type annotations on the left are ordered by significant differences in percentage composition. Differences between populations by cell type are shown in the bar graph on the right. X-axis, horizontal color bars indicate developmental maturity spectrum across cell types as mature, immature, or neuroblasts. Surface fish had a greater proportion of neurons in the excitatory and inhibitory optic tectum cell types (e2: beta-binomial GLM, FDR-adjusted p = 0.001; i5: binomial GLM, FDR-adjusted p = 0.007), as well as in the Raphe (e7: binomial GLM, FDR-adjusted p < 0.001), habenula (e8: beta-binomial GLM, FDR-adjusted p = 0.02), posterior tuberculum projection neurons (e10: binomial GLM, FDR-adjusted p < 0.001), glycinergic hindbrain neurons (HB; i1: beta-binomial GLM, FDR-adjusted p < 0.001), and in cycling neuroblasts (n1: binomial GLM, FDR-adjusted p = 0.0004). Cavefish had a greater proportion of neurons in slow modulatory cerebellar neurons (e5: beta-binomial GLM, FDR-adjusted p < 0.001), cerebellar granule cells (e6: binomial GLM, FDR-adjusted p = 0.014), pineal neurons (e9: beta-binomial GLM, FDR-adjusted p = 0.0025), and immature neuron cell types (im1: beta-binomial GLM, FDR-adjusted p < 0.001; im2: beta-binomial GLM, FDR-adjusted p = 0.0002). **D, E)** Marker gene panels indicating neuron developmental states. **D)** Marker gene panel for neuroblasts, immature, and mature neurons **E)** Marker gene panel for neuroblast subtypes divided by early pre-mitotic progenitors and late-stage, post-mitotic *(elavl3)* progenitors. **F)** Percent composition of neuron developmental stage between populations. Surface fish have a greater proportion of mature neurons (beta-binomial GLM, FDR-adjusted p < 0.001) and neuroblasts (beta-binomial GLM, FDR-adjusted p = 0.0043), while cavefish have a significantly greater proportion of immature neurons (beta-binomial GLM, FDR-adjusted p < 0.001). Error bars in (C) and (F) represent standard error across samples.

### Major cell type annotation and subclustering

Annotations and subsequent data manipulation were performed in R using Seurat. Cell type cluster annotations were based on differentially expressed genes (DEGs) across clusters. This DEG analysis used a pseudobulk approach with sample, rather than cells, as the replicate [44–47]. Pseudobulk analysis was performed by aggregating raw UMI counts across cells within each sample and cell type. Each sample contained snRNAseq count data pooled from approximately 100, 6 dpf larval brains, with a total of four samples each for surface fish and four samples for cavefish. Genes were then filtered within a cell to retain those with at least ten raw counts in at least two samples. The differential expression analysis for DEGs was performed using DESeq2 [48], where counts for each target cell type were compared against all other cell types. Comparisons were performed within each sample and then integrated across samples using a generalized linear model (glm) that included sample identity to account for variability across samples, particularly with respect to *A. mexicanus* population level differences. Briefly, model-fitting employed an empirical Bayes framework, significance was determined using Wald tests with multiple-testing correction, and effect sizes were stabilized using the apeglm estimator for shrinkage of log2 fold change estimates to reduce variance for lowly expressed genes [49]. DEG results for each cluster were then used to assign cell type identities.

Cell type identities were first assigned to three top-level groups: 1) CNS Neurons, 2) CNS Glia, and 3) other non-CNS neuron cell types. We then reclustered cells into these three groups based on marker gene expression, for example, all clusters identified as neurons were grouped together separate from other clusters and then reclustered within this neuron group. In cases where the original clustering produced mixed group clusters, these clusters were further separated into their respective group through subclustering that revealed a single group marker gene profile. If following group reclustering, there was a cluster that clearly belonged to another group, it was reassigned before reclustering anew.

We used an elbow plot to determine the number of PCs to generate reclustered cell types and clustering resolutions were determined using clustering trees generated by the Clustree package with SC3 stability scores [41,42]. For the glia and other non CNS neuron cell types, we ran the FindNeighbors command with harmony reduction, followed by FindClusters and RunUMAP. For the glia subset, we used a PC count of 20 and a clustering resolution of 0.6 and for the other subset, we used a PC count of 30 and a clustering resolution of 0.75. To optimize differences in neuron cell type based on neurotransmitter expression as well as regional identity, we re-ran the PCA and harmony integration prior to the downstream clustering steps. For the neuron subcluster we used 18 PCs and a clustering resolution of 0.8. We further subset the optic tectum cell types from the neuron cluster to examine differences in *A. mexicanus* cell types within a region known to greatly differ between populations. To generate the optic tectum subset, we isolated the relevant clusters, recalculated the variable features, scaling, and PCA prior to clustering with 20 PCs and a resolution of 0.25.

### Differential cell type proportions

To test for differences in cell type proportions between surface and cave populations, we quantified the number of cells in each cell type out of the total cells in that sample. We then fit a binomial glm to each cell type by sample. *A. mexicanus* population and sample sequencing batch were used as covariates, consistent with commonly used approaches in single-cell differential abundance analysis[43–45]. This framework model proportions data on the log-odds scale and treats each pooled sample as an independent biological replicate [46]. The surface population was used as the reference for the cave population, meaning odds ratios greater than one implied a greater proportion in the cave population compared to the surface population. Inference for population effect was based on Wald tests.

Variance between samples for some cell types exceeded that expected under a binomial model (binomial overdispersion > 1.5). In some cases, this variance was explained by batch effects between samples, but in cases where variance was larger than that explained by batch, we fit a beta-binomial model to account for binomial overdispersion [47].This accounted for false positives due to between sample variance. P-values were adjusted across cell type using the Benjamini-Hochberg procedure for false discovery rate.

### DEG testing

Analysis for differentially expressed genes (DEGs) was performed between cell types and between *A. mexicanus* populations. DEG analysis between cell types enabled annotation of each cell type. DEG analysis between surface and cave populations was used to determine population specific expression differences of a single cell type. Analysis used a pseudobulk approach with sample, rather than cells, as the replicate [48–51]. Pseudobulk analysis was performed by aggregating raw UMI counts across cells within each sample and cell type. Each sample contained snRNAseq count data pooled from approximately 100, 6 dpf larval brains, with a total of 4 samples for surface fish and 4 samples for cavefish. Genes were then filtered within a cell to retain those with at least 10 raw counts in at least 2 samples.

Differential expression analysis for DEGs was performed using DESeq2 [52]. For identification of cell type specific markers, counts for eah target cell type were compared against all other cell types, with comparisons performed within each sample and then integrated across samples using a generalized linear model that included sample identity to account for variability across samples, particularly with respect to *A. mexicanus* population level differences. Briefly, model-fitting employed an empirical Bayes framework, significance was determined using Wald tests with multiple-testing correction, and effect sizes were stabilized using the apeglm estimator for shrinkage of log2 fold change estimates to reduce variance for lowly expressed genes [53].

Differential expression analysis for within cell type population comparisons were performed in a similar fashion, with sample pseudobulk aggregation and gene filtration as described above.

DESeq2 analysis used the same Bayes modeling framework and significance testing used to determine differences across cell types, but instead modeled differences between surface and cave populations within a single cell type. Surface fish were used as the reference, so that positive log2fold changes represented higher expression in cavefish relative to surface fish.

In addition to DEG analysis for cell type specific marker genes, we also performed DEG analysis between surface and cave *A. mexicanus* populations within a cell type to determine population specific differences in transcription. Differential expression analysis for within cell type population comparisons was performed in a similar fashion to the DEG analysis for cell type specific marker genes, using the sample pseudobulk aggregation and gene filtration as described above. DESeq2 analysis also used the same Bayes modeling framework and significance testing, but instead modeled differences between surface and cave populations within a single cell type. Surface fish were used as the reference, so that positive log2fold expression changes represent higher expression in cavefish relative to surface fish and negative log2fold expression changes represent higher expression in surface fish relative to cavefish. In cell types with a limited number of cells, e.g. microglia, we also used FindMarkers, with the whole-brain dataset as background for input into the Model-based Analysis of Single-cell Transcriptomics (MAST) package[54].

### Enrichment analysis

The DEGs by cell type and population were tested for signaling pathway enrichment. KEGG enrichment analysis was performed using DAVID bioinformatics [55] for functional annotation with a multiple testing correction of 0.1. All expressed genes detected in our cell type data set served as background against significant DEGs for surface or cave populations.

In cases where DEG number was too limited for significant enrichment, functional groupings for gene expression in that cell type were assigned manually. DEGs were grouped into these functional categories based on well-established gene functions. To avoid selection bias, DEGs were first filtered for significance and a log2fold change cutoff of +/- 0.5, then ranked by significance and effect size, and the top genes selected. Functional categories were then assigned prior to visualization and applied consistently across datasets.

## Results

### Generation of single-cell atlas

To generate a 6 dpf comparative brain atlas in *A. mexicanus,* the brains of 6 dpf surface fish and Pachón cavefish, (∼100 brains/sample) were dissected and flash-frozen. Samples were prepared for snRNA-seq using the 10x Genomics Chromium System, with ∼100 brains per sample. After library alignment and QC filtering (Fig. S1), this yielded 71,251 total cells (29,691 surface and 41,560 cave) with a median of 3,609 unique reads per cell. Median UMI counts did not differ significantly between surface and cavefish samples (Wilcoxon rank-sum test, p-value = 1), indicating high and equivalent quality between populations.

We first divided the cells into three top-level groups: neurons, glia, and other cell types. To divide cells into these major cell type classes, we used 41 transcriptional clusters (Fig. S2A) as a guide to assign cells to one of our three cell type classes based on marker gene expression (Fig. S2B-D). Although many clusters showed homogenous identity and mapped entirely to a single cell class, other clusters contained mixtures of cell classes, such as those that clustered primarily based on regional identity and contained both glia and neurons.

These mixed group clusters were further divided using subclustering in two main ways: 1) by reclustering the initial mixed group cluster and assigning group identities to clearly split subclusters, for example if an initial cluster expressed markers for both neurons and glia, subclustering produced clear divisions into neuron and glia cell types, or 2) when reclustering by group identity yielded a cluster that clearly belonged to a different group, for example reclustering of neuron cell types yielded a clear glia cluster that was reassigned a glia identity before reclustering neurons once more without this cell type. As a result, final cell type groups do not have a one-to-one relationship with initial clustering structure, but nevertheless maintain cell types defined by clustering algorithms within each major group.

The expression of cellular markers was distinct across each major cell type groups, with little overlap between neurons and glia (Fig S2C,D, ST1). These reclustered group subsets, with specific neuronal, glial, and other cell types, allowed us to make comparisons in more specific cell type composition and transcription between surface and cavefish within groups.

### Cell type composition differences between surface and cavefish

To identify compositional and expression differences in specific neuron cell types between *A. mexicanus* populations, we used a combination of conserved cell type markers, validated in *Danio rerio,* for developmental maturity, region, neurotransmitter expression and neuron cell type specificity [56–60](Fig 1A-C). Cell types were identified based on developmental state and divided into progenitor-like neuroblast cells, immature or developing neurons, and fully developed, mature neurons (Fig 1C-E, ST2). Developmentally mature neurons were annotated based on regional markers and were classified by neurotransmitter expression as excitatory (e) or inhibitory (i; Fig. 1C; Fig S3, Fig S4A-E). In some cases, the combination of neurotransmitter, regional and cell type specific markers allowed us to pinpoint a much more specific cell type, such as pineal gland neurons that expressed markers for excitatory signaling (Fig. S3A, B), pineal transcription factors, light sensing, phototransduction, circadian regulation, and melatonin synthesis (Fig. S4E, F).

Differences in cell type composition between surface and cavefish were clearly visible across maturity and brain-region specific markers that co-localized with the pan-neuronal makers *snap25a* and *elavl4* (Fig 1B). Statistical comparisons of percentage composition of cell types between *A. mexicanus* populations revealed differences in 12 of our 24 transcriptionally distinct cell types. These included neuron cell types involved in the regulation of motor circuitry such as the serotonergic Raphe, posterior tuberculum, and glycinergic hindbrain (HB) neurons with greater proportions in the surface fish and slow modulatory cerebellar neurons and cerebellar granule cells with greater proportions in the cavefish (Fig 1C), suggesting broad cellular changes in neural composition across the brain, including those that may reflect adaptive modulation of cavefish motor control systems.

We further examined compositional differences in neurodevelopmental stage between 6 dpf surface and cavefish. We identified two neuroblasts cell types, actively cycling neurogenic progenitors (n1: cycling neuroblasts) and late stage, pre-migratory neurogenic progenitors (n2), two immature neuron cell types (im1, im2), and 20 mature cell types (Fig. 1A-E). To compare neuron development between *A. mexicanus* populations, we grouped cell types into three developmental stages: neuroblasts, immature neurons, and mature neurons, with distinct marker gene expression programs expressed at each developmental stage (Fig. 1D; ST3)

Comparative compositional analysis between *A. mexicanus* populations revealed a significantly greater proportions of neuroblasts and mature neurons in the surface fish and a significantly greater proportion of immature neurons in the cavefish (Fig. 1F). These differences suggest a shift in developmental state allocation at 6 dpf in cavefish that is not consistent with a simple developmental delay, but rather a prolonged maintenance of an immature neuron state. Based on these findings, we conclude there are broad differences in developmental state and cellular composition between surface and cavefish.

Next, we analyzed mature neuron cell types in clusters that defined brain regions and neurotransmitter type (Fig. S3 and S4). Grouping neuron cell types by major brain divisions revealed a greater proportion of cells in the surface fish mesencephalon, consistent with volumetric brain atlas data (Fig S4A, B, ST4; [27,34]). We also grouped neuron cell types by more specific regional marker gene expression within the major brain divisions[56–58]. This further revealed differences in neurons by regions between *A. mexicanus* populations (Fig. S4C, D, ST5). We found a greater proportion of neurons in the posterior tuberculum, optic tectum, and hindbrain of the surface fish and a greater proportion of neurons in the cerebellum and pineal gland of the cavefish (Fig. S4D). The reduction of the cavefish optic tectum and more than 2-fold increase in pineal neuron composition in cavefish is consistent with previously reported neuroanatomical data [26]. Therefore, the snRNA-seq analysis of neuroanatomical cell type composition largely reflects previously published findings from 6 dpf brain atlases and is able to identify defined neuroanatomical regions.

### Comparative analysis of the optic tectum cluster

We next sought to investigate the evolution of cell types within defined brain regions. The optic tectum (OT), the primary visual processing region of the fish brain, is reduced in size in both larvae and adult cavefish, but the cell type composition of this brain region has not been investigated. The optic tectum was identifiable based on the expression of tectal patterning transcription factors that have previously been described in zebrafish (Fig. S3E, F; Fig. S4E). There were two marker gene defined OT cell types from the neuron group recluster, one excitatory cell type (e2) that expressed genes encoding the vesicular glutamate 2 transporter, and one inhibitory cell type (i5) that expressed genes encoding glutamate decarboxylase enzymes and the vesicular GABA transporter (Fig. 1A-C; Fig 2A).

**Figure 2:**
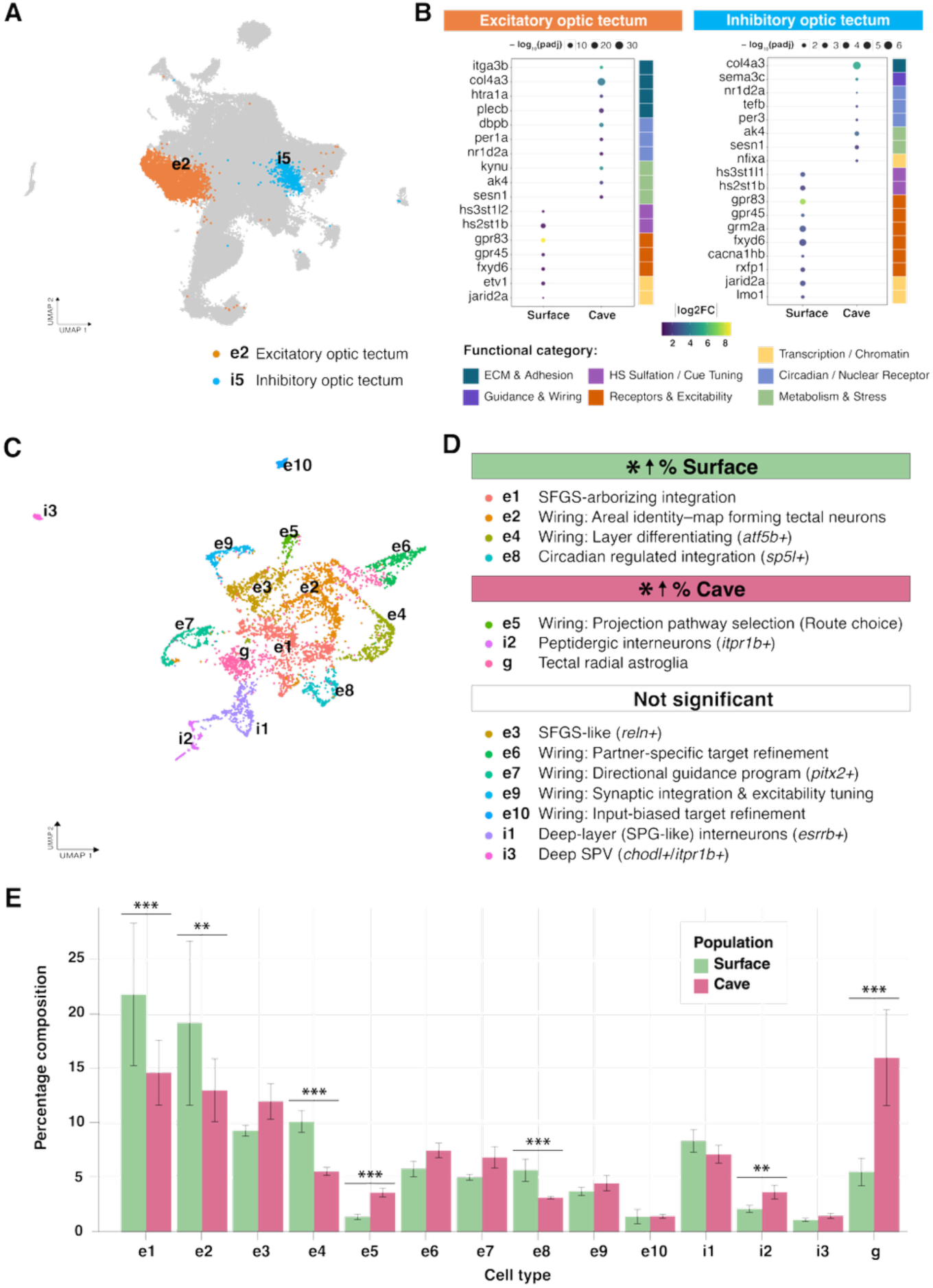
Comparative analysis of optic tectum neuron cell types revealed differences in gene expression and tectal cell type composition between surface and cavefish. **A)** UMAP of CNS neurons highlighting excitatory (e2) and inhibitory (i5) optic tectum neuron cell types. **B)** DEGs for optic tectum cell types between populations with color strips indicating functional group. DEGs distributed into functional groups consistently across excitatory (left) and inhibitory (right) tectal cell types between surface and cave populations. **C-E)** Comparative composition of reclustered tectal cell types. **C)** UMAP of reclustered tectal cell types. **C)** Specific cell type annotations arranged by significant population differences in percent composition. **E)** Percentage composition analysis of tectal cell types between surface and cavefish. Surface fish had a significantly greater proportion of SFGS-arborizing integration neurons (e1; beta-binomial GLM, FDR-adjusted p = 0.0007), circadian regulated integration neurons (e8; beta-binomial GLM, FDR-adjusted p = 0.0006), and two neuron types expressing wiring programs for areal identity (e2; beta-binomial GLM, FDR-adjusted p = 0.0033) and layer differentiating (e4; binomial GLM, FDR-adjusted p < 0.001). Cavefish had a significantly greater proportion of peptidergic interneurons (i2; beta-binomial GLM, FDR-adjusted p = 0.0034), specialized tectal radial astroglia (g; beta-binomial GLM, FDR-adjusted p < 0.001), and neurons expressing a wiring program for route choice (e5; binomial GLM, FDR-adjusted p < 0.001). Error bars represent standard error across samples.

Comparative analysis of differentially expressed genes within these two optic tectum cell types revealed divergence in functional programs (Fig. 2A,B, ST6). Functional divergence in gene expression between surface fish and cavefish was consistent across both cell types (Fig 2B). Surface fish tectal neurons display upregulated expression of genes involved in input signal modulation, extracellular cue tuning (heparan sulfate modification), and upstream gene regulatory factors (transcription/chromatin). Cavefish tectal neurons display upregulated expression of genes involved in state-dependent transcriptional regulation (circadian/nuclear receptor), extracellular structural interface (ECM and adhesion), cellular maintenance (metabolism and stress), and a single different transcriptional regulator (*nfixa*). This is consistent with broad changes in neural activity in the cavefish tectum associated with loss of light-sensing [61].

To investigate differences in cell type within the tectum, we combined our tectal neuron cell types and reclustered into ten excitatory cell types, three inhibitory cell types, and one tectal specific glial cell type (Fig. 2C, D, ST7). We characterized our new set of tectal cell types by their neurotransmitter expression (Fig. S5A) and top differentially expressed genes (Fig. S5B). Many of the top cell type DEGs indicated expression differences associated with differences in synaptic development and connectivity (Fig. S5C; ST8). Compositional analysis of tectal cell types revealed maturing tectal cell type wiring programs enriched for tectal patterning in surface fish and an enrichment for route choice in the cavefish. In addition to wiring program differences, there were also compositional differences in more mature cell types, including an increased proportion of stratum fibrosum et griseum superficiale (SFGS) arborizing integration neurons and circadian regulated integration neurons in the surface fish and peptidergic interneurons and tectal radial astroglia in the cavefish (Fig. 2D, E). Therefore, the cell type composition and gene expression within cell types of the optic tectum has diverged between surface fish and cavefish and may reflect tectal circuit reorganization in the absence of light induced neuron development.

### Comparative analysis of pineal clusters

The loss of visual input is also associated with changes in the light-sensing function of the pineal gland that remains functional in cavefish [62]. In mammals pineal neurons harbor an endogenous circadian clock that contributes to rhythmic synthesis and secretion of melatonin [63]. Comparison of relative clock gene expression in pineal neurons revealed divergence in the expression of positive and negative clock transcription components between *A. mexicanus* populations. Surface fish showed elevated expression levels of genes in the positive feedback, while cavefish showed elevated expression levels of the negative feedback (Fig. 3B-F). Although limited to a single timepoint, this difference in expression in clock gene transcription indicates changes in clock-gene regulation within the pineal gland.

**Figure 3.**
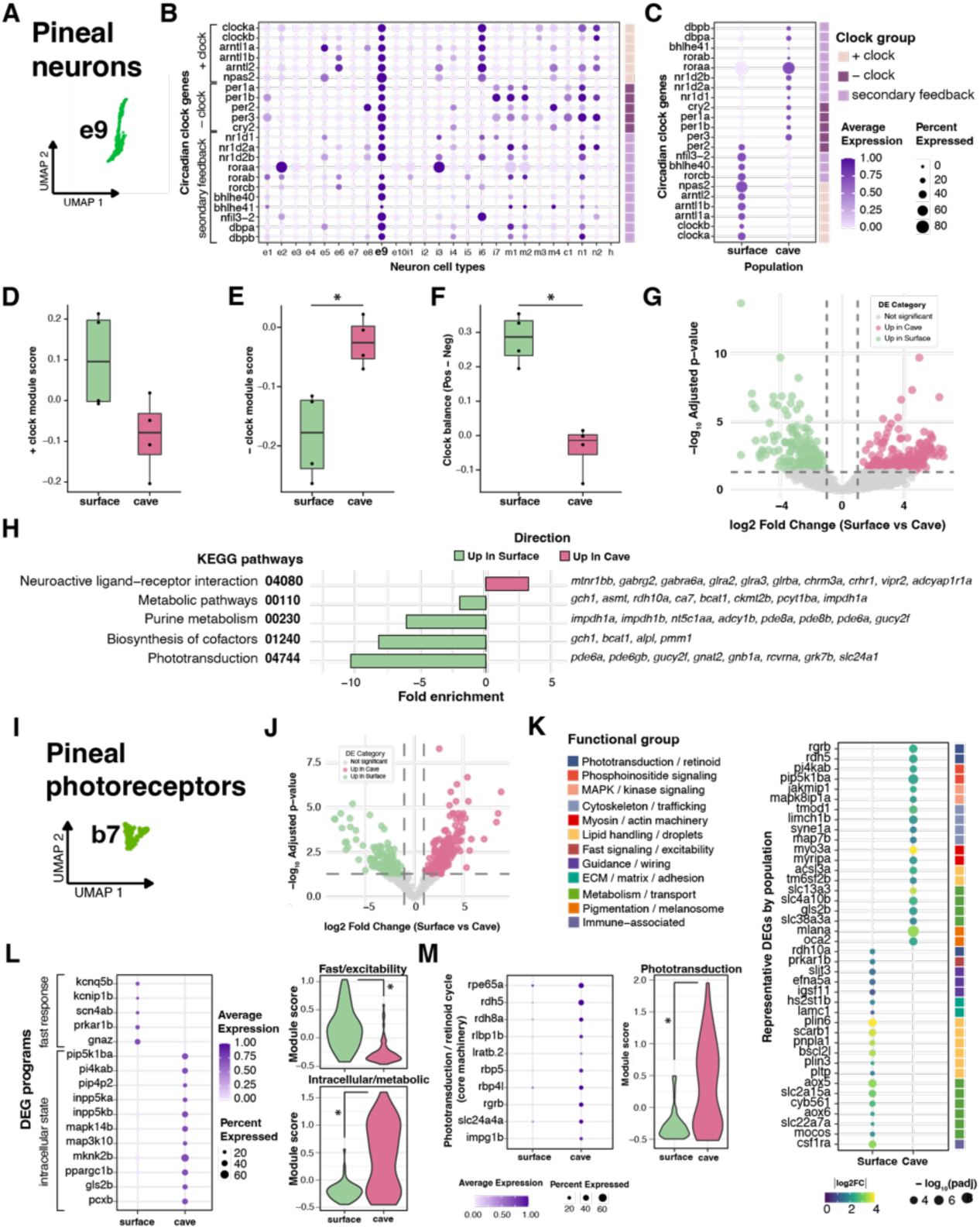
Gene expression differences between *A. mexicanus* populations for pineal gland cell types. Gene expression differences for pineal neurons (A-H) and pineal photoreceptor cells (I-M), a specialized non-canonical neuron subtype. **A)** UMAP distribution of pineal neurons (e9) subset from CNS neurons. **B)** Marker panel of circadian clock genes showing predominant expression in pineal neurons compared to other neuron cell types. Genes are grouped as positive, negative, and secondary feedback regulatory components of the clock system. Clock components are also labeled in the color bar at the right as reference. **C)** Comparative expression of circadian clock genes between populations revealed divergence in the expression of positive and negative clock transcription components. Color bar refers to clock component divisions labeled in (B). **D, E)** Module expression scores by sample for positive (D) and negative (E) clock components. Cave fish had significantly higher expression of negative clock components by sample compared to surface fish (Wilcoxon: p-value = 0.029). **G)** Clock balance score calculated as the per sample difference between positive and negative clock scores by population. There was a significant per sample difference in clock balance score between surface and cave populations (Wilcoxon: p-value = 0.029). **G, H)** DEG analysis between populations for pineal neurons. **G)** Volcano plot showing distribution of pineal neuron DEGs between surface and cave fish. **H)** Significant KEGG pathway enrichment (FDR < 0.1) for pineal neuron DEGs showed surface enrichment for phototransduction and related mechanisms and cavefish enrichment for neuroactive ligand-receptor interaction. KEGG pathway and ID are listed on the left and genes driving enrichment are listed on the right. **I)** UMAP distribution of pineal photoreceptors (b7) subset from other non-CNS neurons. **J-M)** Differential gene expression for pineal photoreceptors between *A. mexicanus* populations. **J)** Volcano plot distribution of pineal photoreceptor DEGs between surface and cavefish. **K)** Representative pineal photoreceptor DEGs with color strip highlighting functional DEG groups across *A. mexicanus* populations. **L)** Differential expression of genes related to fast response and intracellular state programs. Gene panel and associated module scores revealed increased gene expression in surface fish pineal photoreceptors for fast-response/excitability programs (Wilcoxon: p-value = 0.0304) and increased gene expression in cavefish for intracellular state modulatory programs (Wilcoxon: p-value = 0.0304). **M)** Gene panel and associated module scores revealed increased gene expression in cavefish for phototransduction and retinoid cycle core machinery (Wilcoxon: p-value = 0.0304).

To determine if the identified differences in the gene expression of pineal neurons correlated with functional changes, we performed functional enrichment of differentially expressed genes in pineal neurons between *A. mexicanus* populations. We used pseudo-bulk, sample level analysis to determine the transcriptomic differences in expression within pineal neurons between surface and cavefish (Fig. 3G, ST9). KEGG enrichment analysis of these DEGs revealed differences between surface and cave populations following multiple testing correction (FDR ≤ 0.1; Fig. 3H; ST10). Surface fish showed enrichment for phototransduction, driven by canonical components of the phototransduction. Surface fish were also enriched for purine metabolism, metabolic pathways, and biosynthesis of cofactors, all pathways that reflect the metabolic demands and cyclic nucleotide turnover associated with the phototransduction process (Fig 3J). In contrast, cavefish pineal neurons showed enrichment for neuroactive ligand-receptor interaction, with genes that code for a broad array of neurotransmitter and neuromodulatory receptors, including those for GABA, glycine, glutamate, and ACh. This increase in input modulation raises the possibility of major changes in the regulation of pineal neurons between surface and cavefish.

### Comparative analysis of non-neuronal or glial cell types

We next sought to define the identity of cells that do not transcriptionally classify as neurons or glia from the CNS. Subclustering of these cells yielded 25 distinct cell types, seven of which represented previously identified CNS brain cell types (b) and 18 of which represented peripheral tissue contamination from tissue dissection (Fig. S6A, B, ST10). Compositional analysis of the CNS brain cell types revealed a greater proportion of meningeal fibroblasts and choroid plexus epithelial cells in surface fish and a significantly greater proportion of blood brain barrier (BBB) endothelial cells and meningeal lymphatic endothelial cells (mLECs) in cavefish (Fig. S6C).

Within this subset of other non-canonical CNS cell types we identified a group of pineal photoreceptors cells distinct from the pineal neuron cell type (Fig. 3I). Pineal photoreceptor cells are specialized sensory neurons embedded in the pineal epithelium that do not express typical neuronal markers or neuronal morphology but instead show similar molecular characteristics to retinal-pigment epithelial cells [64]. We identified these cells by the expression of pineal photoreceptor marker genes and the lack of expression of retinal marker genes (Fig. S6D) [65–68]. Analysis of population DEGs in pineal photoreceptors between surface and cavefish (Figure 3J, ST11) showed a divergence in functional programs (Figure 3K). Pineal photoreceptors in surface fish displayed increased expression of genes associated with fast electrical responsiveness (fast signaling/excitability), membrane lipid storage and turnover (lipid handling/droplets), and substrate transport processes (metabolism/transport). In cavefish, pineal photreceptors displayed increased expression of upregulated genes including those involved in chromophore regeneration and photoreceptor maintenance machinery (phototransduction/retinoid), regulation of membrane phospholipid signaling and second messenger production (phosphoinositide signaling), intracellular signaling cascades (MAPK/kinase signaling), as well as cytoskeletal/trafficking regulators and pigmentation-associated components.

Gene set analysis of pineal photoreceptor DEGs between *A. mexicanus* populations further identified program level differences between surface and cavefish. Genes associated with fast response, intracellular state programs, and phototransduction maintenance were differentially expressed, both in average expression and in gene set module scoring, between populations (Fig. 3L, M). Surface pineal photoreceptors showed increased expression for fast response programs, whereas cave pineal photoreceptors showed increased expression for intracellular state regulators (Figure 3L). Cavefish also showed increased expression for phototransduction genes associated with the retinoid-cycle and chromophore regeneration (Figure 3M). These findings raise the possibility of functional differences in the deep-brain pineal photoreceptors.

### Analysis of glial cells

To date, the molecular identity of glia in *A. mexicanus* and evolved changes between surface and cavefish has not been investigated. To identify specific glial subtypes, we reclustered the cells differentially expressing canonical glia markers from the larger whole-brain dataset and subset into a glia only group for comparison across glial cell types. We identified radial glia, astrocytes, tanycytes, ependymal cells, oligodendrocytes and their precursors, and early microglia cells with myeloid lineage characteristics based on previously described zebrafish markers (Fig. 4A; Fig. S7A, ST13). Initial clustering yielded five radial glia cell types, differentiated based on regional or cell type specific markers that indicated ventricular zone niche location (Fig. S7B, C, ST14). Compositional differences across subtypes of glia revealed that cavefish have a higher proportional composition of astrocytes, tanycytes, and microglia, as well as radial glia from the cerebellar/isthmic (Fig. 4B; Fig. S7D). Therefore, there are broad differences in glial subtypes in cavefish that likely reflect differences in developmental allocation and neuron regulation between *A. mexicanus* populations.

**Figure 4:**
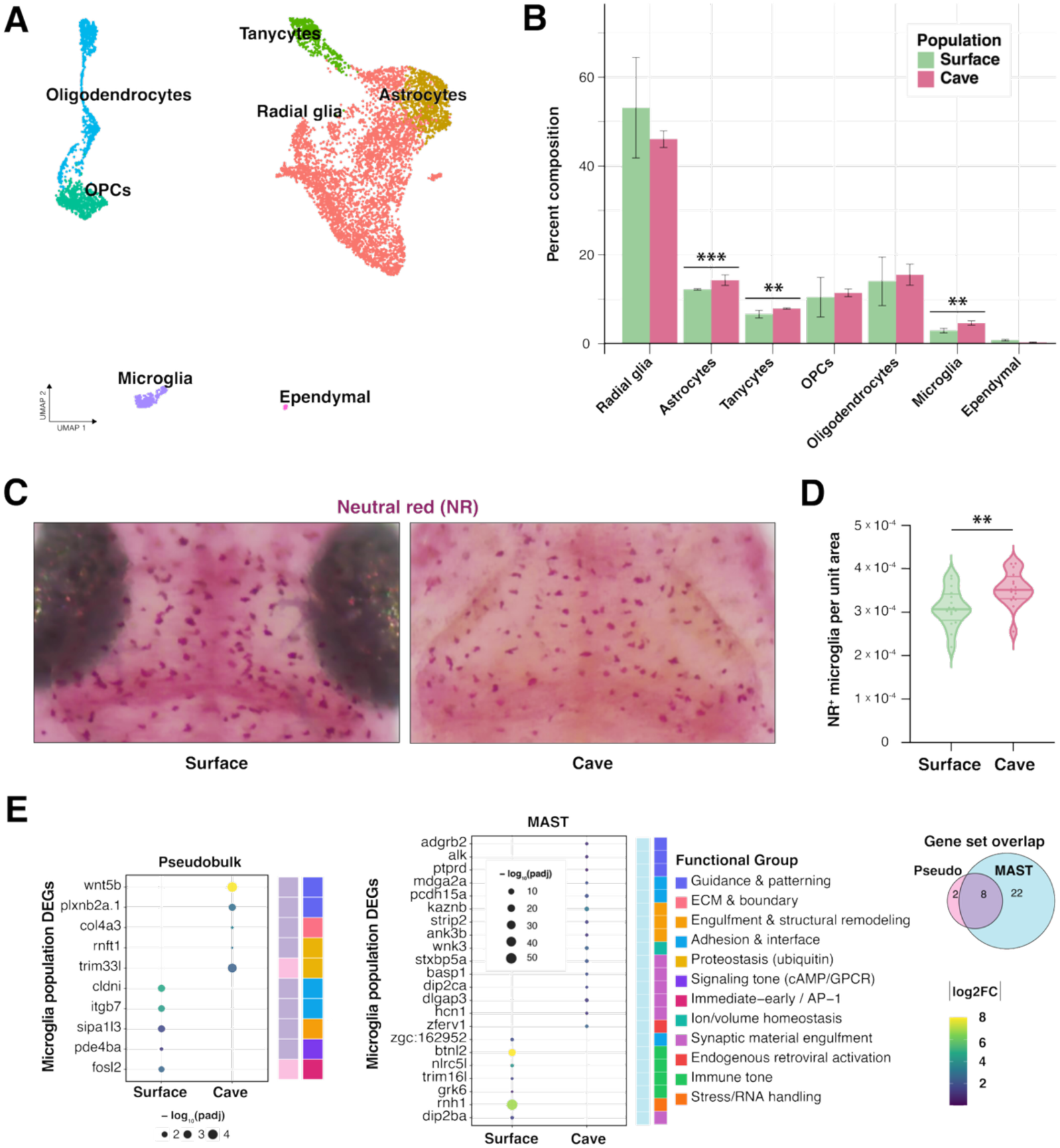
Glial cell type differences between *A. mexicanus* populations. **A)** UMAP of 7 main glial cell types. **B)** Percent composition of glial cell types for surface and cave populations. Cavefish had a significantly greater percentage composition of astrocytes, (binomial GLM, FDR-adjusted p = 0.0003), tanycytes (binomial GLM, FDR-adjusted p = 0.001), and microglia (beta-binomial GLM, FDR-adjusted p < 0.001). **C)** Neutral red staining and quantification of microglia. **D)** Relative number of microglia is higher in cavefish (p<0.001). **E)** Microglia DEG analysis expression panels by population calculated at the sample level (left) and the cell level (right) with color strips showing gene enrichment overlap between analysis levels and differential microglia functional enrichment between populations.

Microglia are not only critical regulators of phagocytosis debris clearance, but are also regulated by environmental processes including sleep and immune response [69,70]. To validate the identified differences in microglia identified with sn-RNAseq we labeled microglia with the lysosomal marker neutral red in 6 dpf surface and cavefish. The number of microglia were significantly greater in cavefish than surface fish, verifying compositional snRNAseq analysis (Fig 4C). To characterize transcriptional changes in microglia between *A. mexicanus* populations in a cell type with a small number of cells (244 total, 98 surface and 146 cave), we performed DEG analysis at both the sample level and the cell level, using pseudobulk with DESeq2 (ST15) and FindMarkers with MAST (ST16), respectively, and grouped gene enrichment by function (Fig. 4D). Surface fish microglia showed upregulated expression of genes involved in immune surveillance and responsiveness including adhesion and barrier associated genes, regulators of GPCR/cAMP signaling, as well as cytoskeleton and morphology genes. Cavefish microglia upregulated genes involved in structural integration and homeostatic maintenance including developmental guidance and patterning programs, extracellular matrix and boundary-associated components, and proteostasis-related regulators. Cell-level analysis further resolved differences in microglia between populations. Surface microglia increased expression in genes related to immune tone regulation and stress-associated RNA handling. Cavefish microglia increased expression in genes related to synaptic material engulfment and engulfment-associated cytoskeleton remodeling. Together, these changes raise the possibility that surface fish microglia are primarily in a surveillance state, primed to mediate an immune response and regulate the degree of resulting inflammation, whereas cavefish microglia have shifted to a more activated state, with a greater role in debris clearance and synaptic pruning.

## Discussion

Here, we present the first comparative single-nucleus RNA-sequencing (snRNA-seq) atlas of the developing *Astyanax mexicanus* brain. In zebrafish, comprehensive brain atlases have provided detailed resolution of neural cell types and developmental trajectories [29,32,33]. We used established markers for zebrafish neuronal and glial subtypes to robustly annotate cell populations in *A. mexicanus* [31,71]. These results enable rigorous comparisons between surface and cave morphs and, importantly, allow direct integration with existing zebrafish datasets, positioning *A. mexicanus* as a powerful comparative system for studying brain evolution and development.

Single-cell sequencing is a powerful approach for identifying cell type–specific changes within comparative models of evolution. A previous single-cell atlas comparing the adult hypothalamus of zebrafish, surface *Astyanax mexicanus*, and Pachón cavefish revealed both conserved neuronal populations and lineage-specific transcriptional specializations associated with behavioral evolution. Narrowing the focus to a single region of the adult brain allowed for quantification of differences in defined classes of neuropeptides, including increases in sleep and orexigenic neuropeptides [30]. In addition, comparative single-cell atlases of cichlid brains have identified diversification in neuronal and glial transcriptional programs associated with species-specific behaviors and ecological niches [29]. Together, these studies highlight how single-cell approaches can uncover both conserved cellular architectures and evolutionary innovations across vertebrate brains. As single-cell atlases are increasingly applied across populations of the same species or closely related species, they provide an opportunity to move beyond traditional comparisons of gene expression or gross neuroanatomy and instead investigate how the evolution of specific cell types, transcriptional programs, and cellular compositions contributes to changes in neural circuit function and behavior.

Cavefish have evolved numerous neurobiological differences, including an expansion of dopamine- and hypocretin-expressing neurons, suggesting widespread remodeling of neural circuits that modulate behavior [27,72,73]. Although some cell types were difficult to detect, presumably due to their relatively low abundance, we were nevertheless able to identify differences in cell type composition across the brain. These include shifts in neuron developmental allocation, differential enrichment of neuron cell types that regulate motor systems, and compositional increases in several glial cell types compared to surface fish.

Further, many cell types and brain regions hypothesized to contribute to cave-associated behaviors remain poorly understood at the molecular level. For example, the pineal gland has been shown to regulate an escape response in cavefish, whereas the optic tectum has been proposed to be repurposed for non-visual processing [74,75]. Our analysis identifies broad shifts in the cell type composition that is predicted to associate with defined neuroanatomical regions, as well as differential transcriptional regulation, providing new avenues for investigating the function of diverse brain cell types and their contribution to evolved behaviors.

Glial cells are critical regulators of brain function, including neurodevelopment, behavior, and synaptic physiology. Evolved differences in glial function have been shown to contribute to behavioral evolution, including changes in sleep regulation and neural circuit activity. Despite extensive changes in neural function, behavior, and neuroanatomy, the role of glia in the evolution of brain function in *A. mexicanus* had not previously been investigated. Here, we identified a proportional increase in tanycytes, astrocytes, and microglia in a cave morph of *A. mexicanus*, suggesting broad differences in glial cell composition and function. These glial cell types have been linked to processes relevant to cave-associated phenotypes. For example, tanycytes localize to the hypothalamus and have been shown to regulate sleep in zebrafish [76]. Astrocytes broadly regulate synaptic transmission and neurotransmitter release, suggesting a potential role in shaping neural circuit physiology [77]. In zebrafish, microglia clear cellular debris including amyloid-β from the brain, and this process has been shown to be sleep-dependent in mammals [78,79]. Together, these findings suggest that evolutionary changes in glial cell composition and function contribute to the neural and behavioral adaptations observed in cavefish.

Here, we chose to develop an atlas in larval fish at 6 dpf. In both zebrafish, and *A. mexicanus*, this stage has been widely used for behavioral neuroscience experiments because the brain is transparent, allowing for whole-brain imaging. In *A. mexicanus,* cave populations have evolved differences in sleep and locomotor regulation, startle response, prey-capture, and olfaction [80–83]. In addition, multiple brain atlases at this stage provide detailed resolution of differences in neuroanatomy [26,27,84]. The molecular-based cell type map generated in this data set provides the opportunity to identify potential changes in function. This data set also provides the opportunity for future studies using spatial-single cell sequencing that will combine the power of the single-cell atlas presented here and existing neuroanatomical brain atlases [85].

The development of this single-cell atlas provides measurements of differential gene expression across cell types within the brain. The comparative differences in gene expression can provide novel insights into function, including differential regulation of clock-genes within the pineal gland, a brain region associated with regulation of circadian behavior and sleep. The identification of these genes also provides the opportunity to label defined cell types using transgenic Tol2 promoters that have been implemented in *A. mexicanus*. In addition, the application in CRISPR-Cas9 that has been used in *A. mexicanus,* and screening in F0 fish that has been optimized in zebrafish, should allow for functional validation of genes [86]. These approaches could be applied to validate the functional role of genes to test for novel regulators of behavior.

Together, this work establishes a foundational molecular atlas of the developing *Astyanax mexicanus* brain and provides a framework for investigating how neural cell types evolve to support behavioral adaptation. Although this study specifically compares surface fish and the Pachón population of cavefish, behavioral and neuroanatomical differences are also widespread in Molino and Tinaja populations, providing potential for future work expanding brain atlases to additional cave populations [27]. By integrating single-nucleus transcriptomics with comparative analyses between surface and cave morphs, our study reveals shifts in neuronal and glial composition and gene expression programs that may contribute to evolved differences in sleep, sensory processing, and locomotor behavior. The conservation of major teleost cell types enables direct comparison with zebrafish resources, while the identification of cave-associated transcriptional changes highlights candidate genes and cell populations for functional study. The data from this single-nucleus cell atlas can be combined with emerging genetic and imaging tools in *A. mexicanus*, including transgenesis, CRISPR-based perturbations, and whole-brain functional imaging to validate gene function [61,87,88]. Together, this atlas provides a resource for linking evolutionary changes in gene regulation and cell type composition to neural circuit function and behavior.

## Data Availability

Raw snRNA-seq reads and processed cell-count matrices are available in NCBI’s Gene Expression Omnibus (GEO) database under accession GSE328673.

## Acknowledgements

The authors are grateful to Lily Pena (Texas A&M) for technical support. This work was supported by NIH award R240D030214 to ACK and WW, BSF 2021-177 to ACK; NIH (R15-MH132057) and the BSF (2019-262) to ERD.

---

**Supplementary Figure 1:**
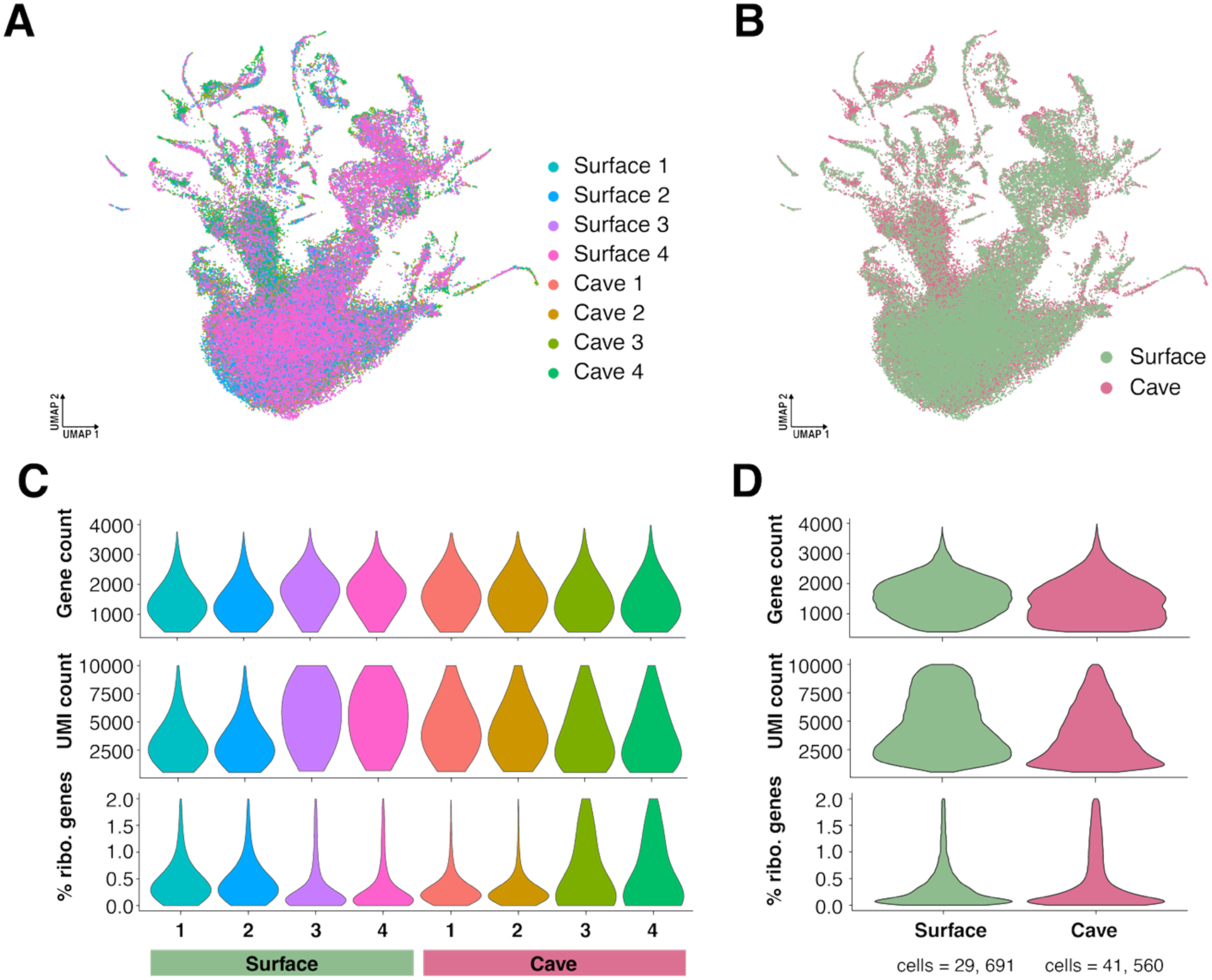
snRNA-seq sample integration and quality control. **A-B)** Cells were integrated using harmony across surface and cave *A. mexicanus* samples. **A)** UMAP showing integration by sample. **B)** UMAP showing integration by *A. mexicanus* populations. C-D) Quality control (QC) thresholds for gene count (< 40,000 genes), UMI count (< 20,000 UMIs), and percent ribosomal genes (< 2% ribo. genes). **C)** QC thresholds by sample yielded 71,251 cells that were consistent across samples. **D)** QC thresholds by *A. mexicanus* yielded 29,691 surface fish cells and 41,560 cavefish cells. There were no significant differences between populations.

**Supplementary Figure 2:**
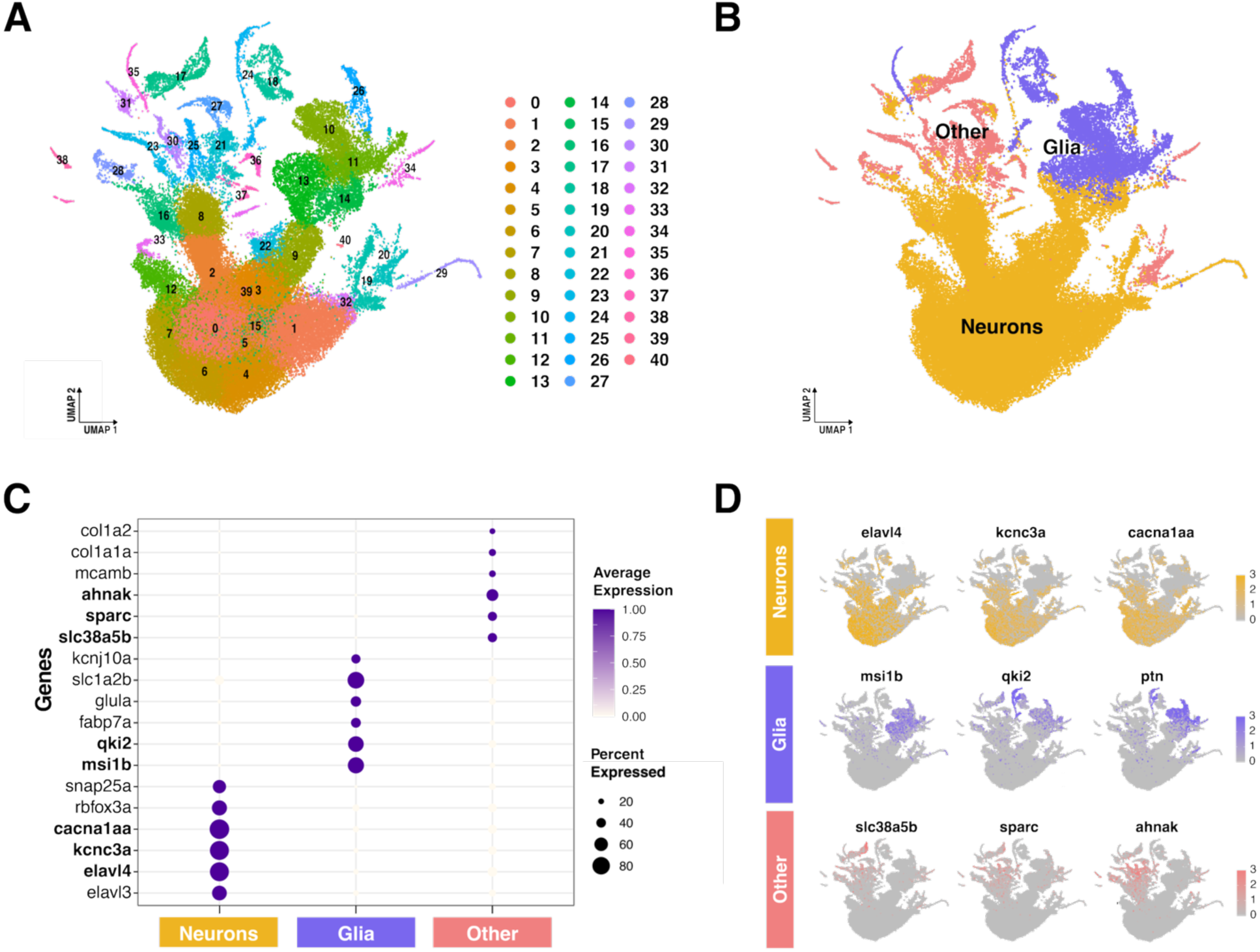
snRNA-seq annotation of Larval *A. mexicanus* major cell type groups. **A)** UMAP of initial 41 transcriptional cell type clusters used for major cell group annotation. **B)** UMAP of annotated cell type groups: CNS neurons, CNS glia, and all other cell types. **C)** Representative marker gene panel for cell type groups. **D)** Marker gene distribution of cell type groups on UMAP.

**Supplementary figure 3:**
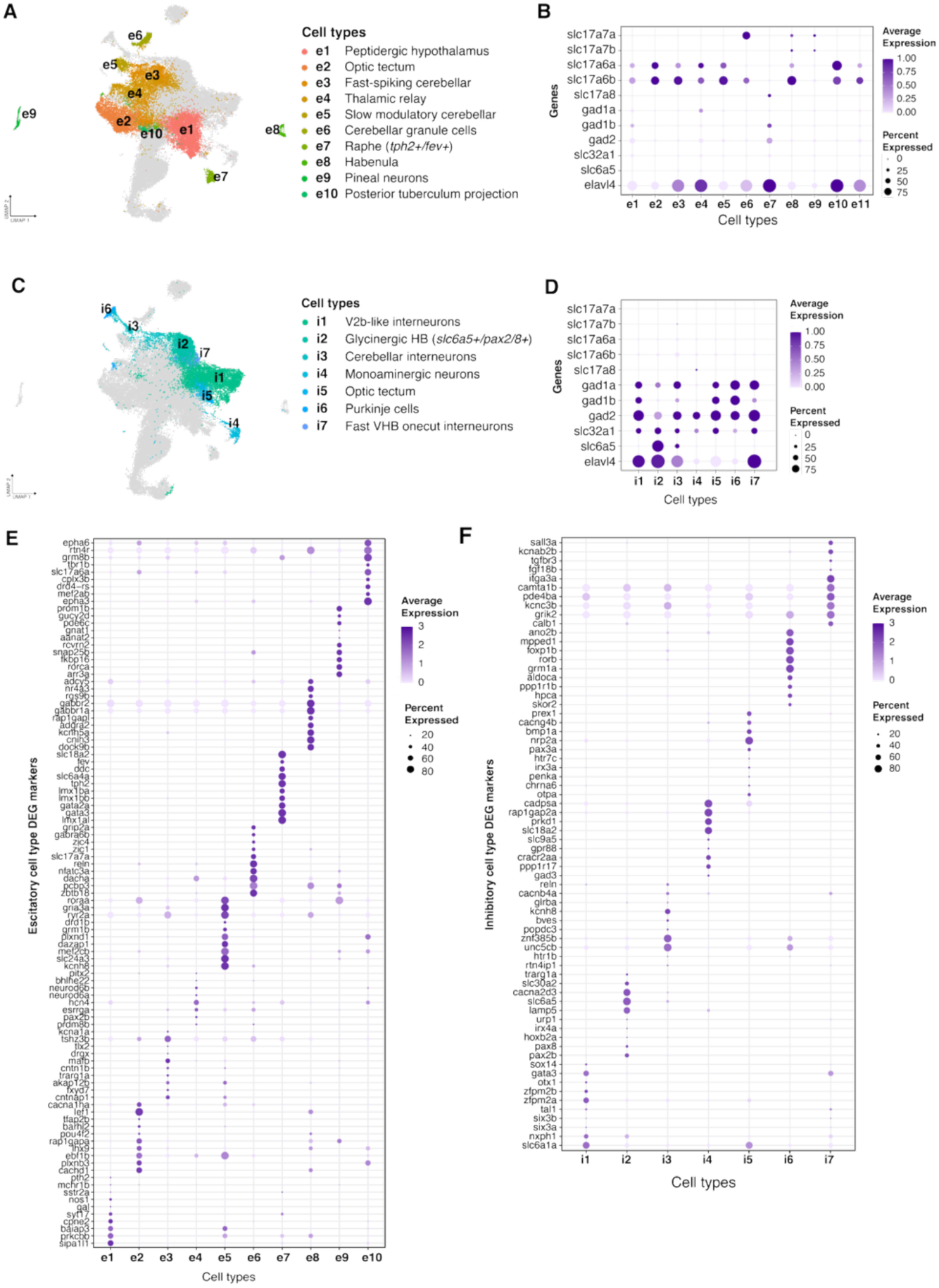
Neuron cell type annotations by excitatory or inhibitory neurotransmitter expression. **A)** Neuron UMAP highlighting excitatory cell types with annotation legend. **B)** Dot plot showing expression of excitatory and neuronal (*elavl4/HuC*) marker genes. *Slcl17a* genes for vesicular glutamate transporters were used as markers for excitatory glutamate neurotransmission **C)** Neuron UMAP highlighting inhibitory cell types with annotation legend. **D)** Dot plot showing expression of inhibitory and neuronal marker genes. Inhibitory marker genes for GABA and glycine neurotransmission included *gad* genes for glutamate decarboxylase enzymes, *slc32a1* for vesicular GABA transporter, and *slc6a5* for glycine transporter 2. **E)** Marker gene panel by excitatory cell type. **F)** Marker gene panel by inhibitory cell type.

**Supplementary figure 4:**
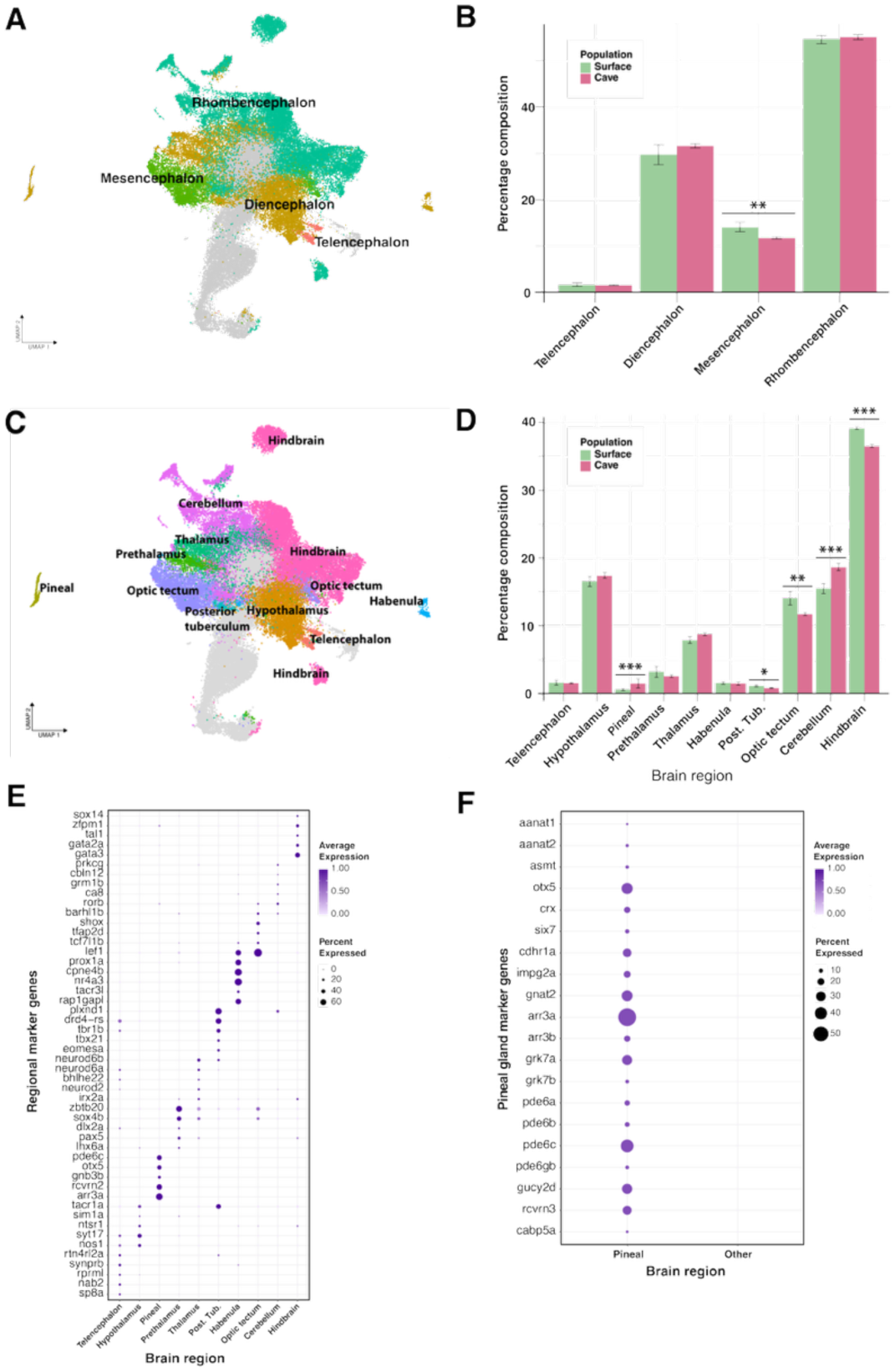
CNS neuron composition varies between *A. mexicanus* populations by major brain division and by brain region. **A)** UMAP showing mature cell types merged by brain divisions. **B)** Comparative composition of brain divisions between surface and cavefish. Surface fish have a significantly greater percentage composition of neurons in the mesencephalon compared to cavefish (beta-binomial GLM, FDR-adjusted p = 0.0021). **C)** UMAP showing mature cell types merged by brain region. **D)** Comparative percentage composition of brain regions between surface and cavefish. Surface fish had a significantly greater percentage composition of neurons in the posterior tuberculum (post. tub.; beta-binomial GLM, FDR-adjusted p = 0.0486), the optic tectum (beta-binomial GLM, FDR-adjusted p = 0.0021), and the hindbrain (binomial GLM, FDR-adjusted p < 0.0001), while cavefish had a significantly greater percentage composition of neurons in the pineal gland (beta-binomial GLM, FDR-adjusted p = 0.0001) and the cerebellum (binomial GLM, FDR-adjusted p < 0.0001). **E)** Marker gene panel by brain regions. **F)** Specific marker gene panel for pineal neurons. Error bars in (B) and (D) represent standard error across samples.

**Supplementary figure 5:**
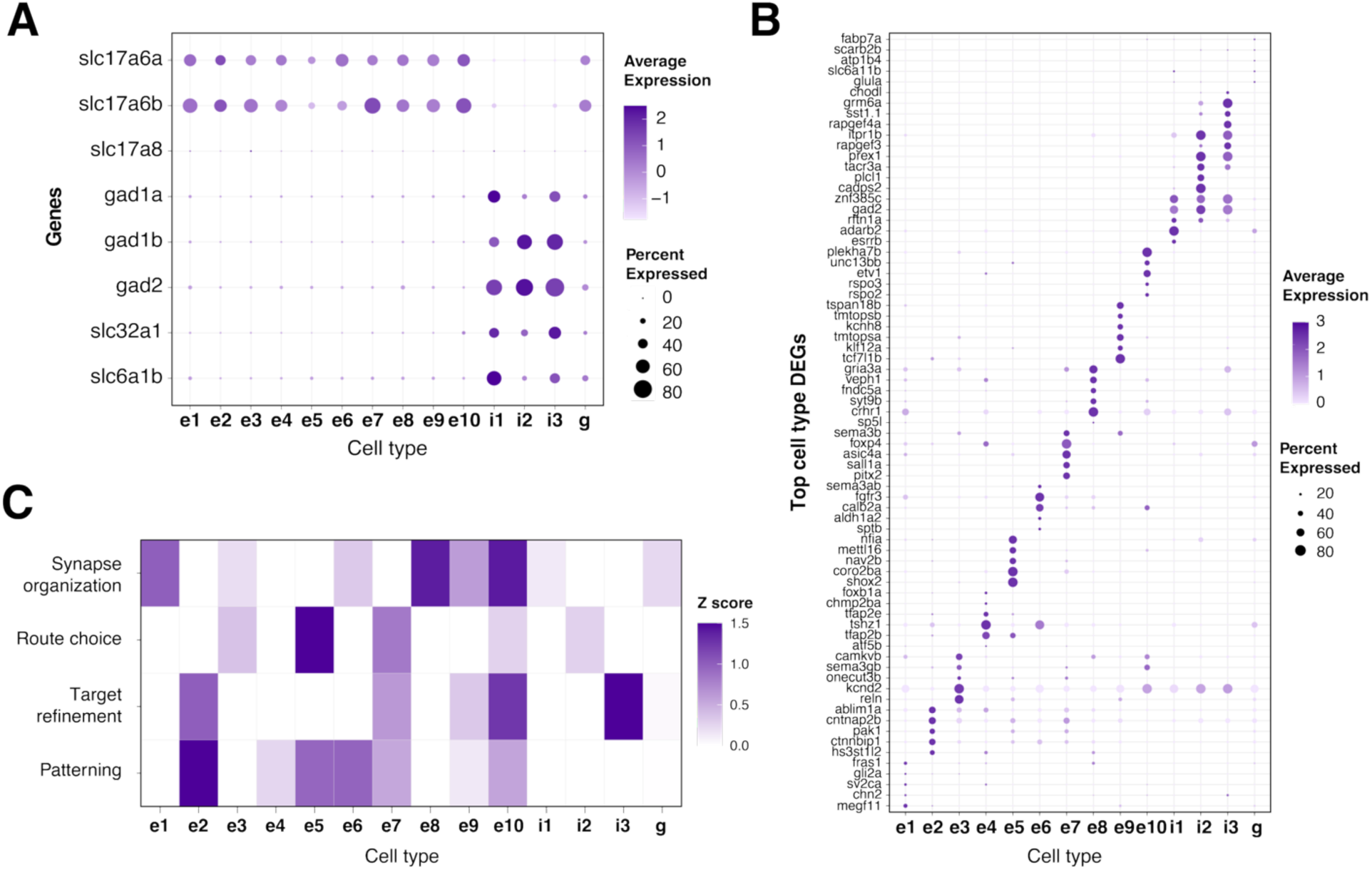
Marker gene and wiring program expression in optic tectum specific cell types. **A)** Dot plot showing expression of excitatory (e) and inhibitory (i) gene expression across reclustered tectal cell types. **B)** Marker panel for tectal cell types. **C)** Expression of wiring programs across cell types. Gene sets used to calculate module scores representing patterning, axon guidance (route), target refinement, and synaptic organization programs are provided in Supplementary Table 8. Heatmap values represent median module scores per cell type, z-scored across cell types.

**Supplementary figure 6:**
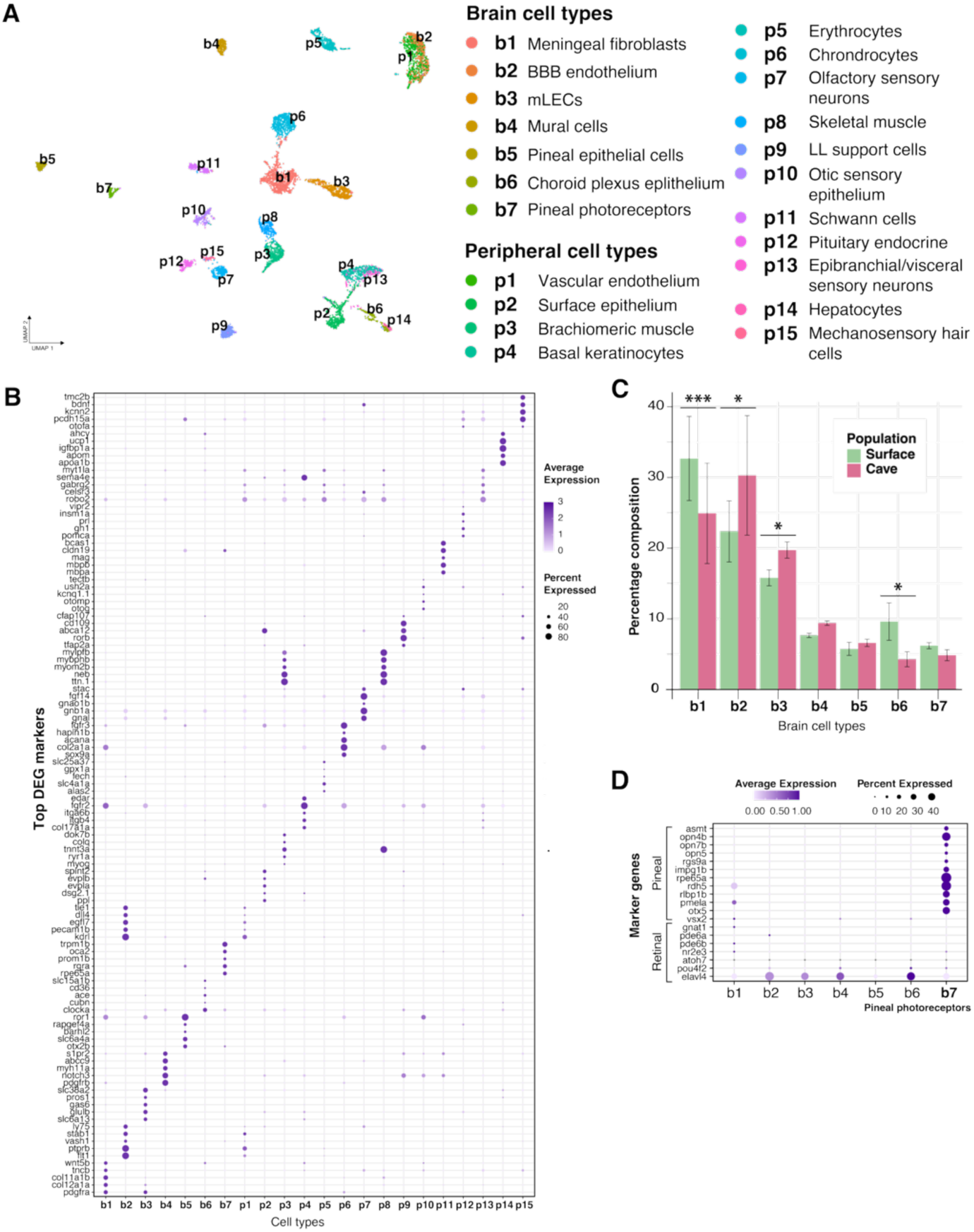
Comparative analysis of other non-CNS neuron and non-glial cell types revealed differences in percentage composition between *A. mexicanus* populations. **A)** Cell types that did not belong to CNS neurons or glia major cell type groups were pooled and annotated as either brain cell types (b) or peripheral cell types (p). These “other” cell types are displayed as a UMAP with specific annotation labels in the adjacent legend. **B)** Marker gene panel of top DEGs by other non-CNS neuron or glia cell types. **C)** Comparative percentage composition of “other” brain cell types. Surface fish had a significantly greater percentage composition of meningeal fibroblasts (b1: binomial GLM, FDR-adjusted p = 0.0005) and choroid plexus epithelial cells (b6: beta-binomial GLM, FDR-adjusted p = 0.0481), while cavefish had a significantly greater percentage composition of blood-brain-barrier (BBB) endothelial cells (b2: beta-binomial GLM, FDR-adjusted p = 0.015) and meningeal lymphatic endothelial cells (mLECs; b3: binomial GLM, FDR-adjusted p = 0.0111). Error bars represent standard error across samples. **D)** Expression of pineal photoreceptor marker genes in b7 with distinct lack of retinal marker genes expression, verifying a pineal rather than retinal identity for this cluster.

**Supplementary figure 7:**
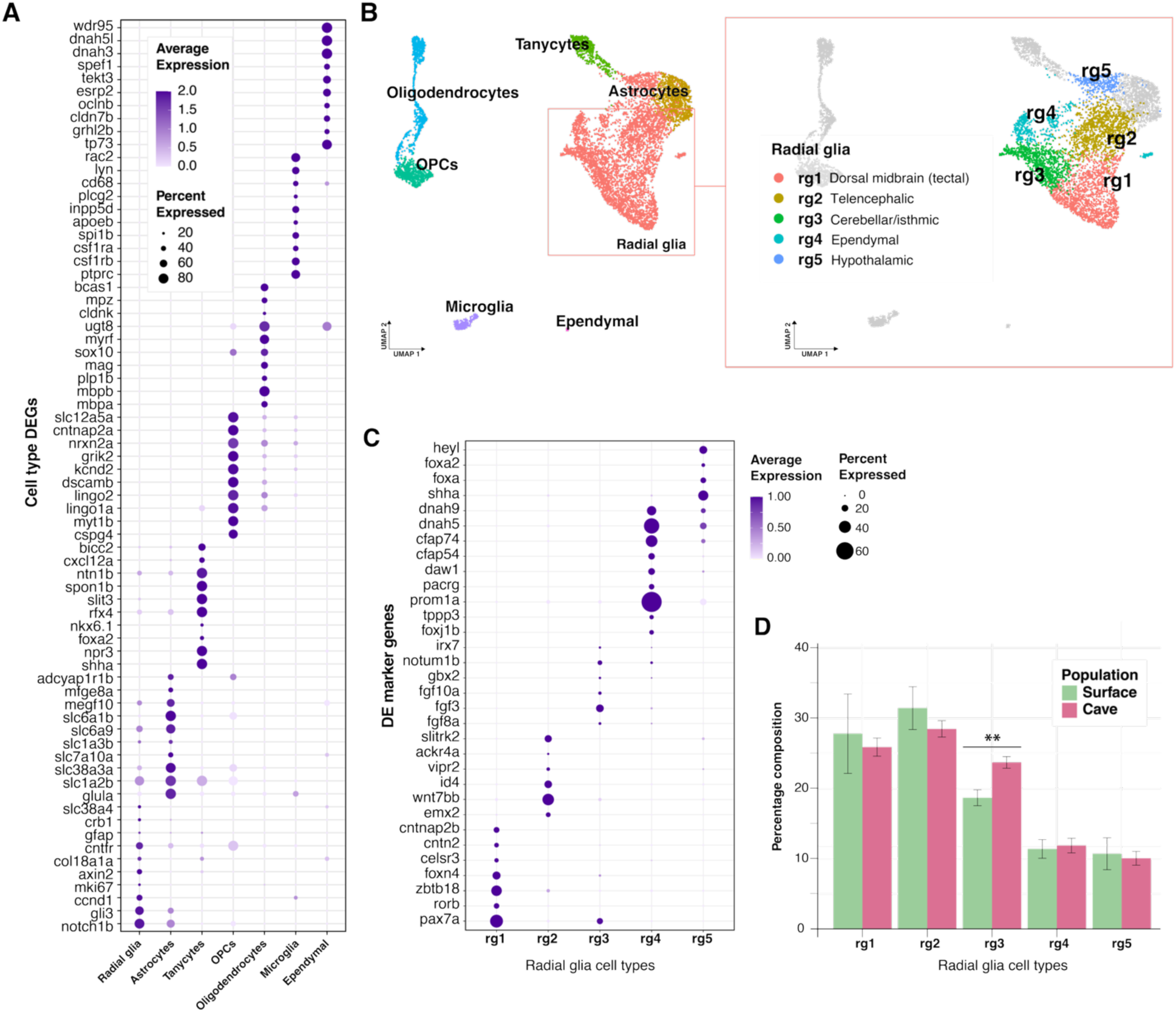
Comparative analysis of radial glia cell types revealed differences in percent composition between surface and cavefish. **A)** Marker gene panel across glial cell types. **B)** UMAPs of glial cell types. Main glial cell types shown in the left UMAP and specific radial glial cell types highlighted on the right with annotation legend. **C)** Marker gene panel across radial glia cell types. **D)** Percent compositional analysis of radial glia cell types between surface and cave populations revealed a greater proportion of radial glial cells from the cerebellar/isthmic region in the cavefish (binomial GLM, FDR-adjusted p = 0.0036). Error bars represent standard error across samples.

## Supplemental table legends

**Supplementary table 1: Differentially expressed genes (DEGs) for major cell type groups.** DEGs were calculated using a pseudobulk deseq2 approach that compared gene expression of each cell type group (neurons, glia, and other non-neuronal/glia cell types) to all other cell type groups (see Methods). The table reports adjusted p-values (padj) using BH correction for FDR, log2 fold change (log2FoldChange), standard error (lfcSE), raw p-values (pvalue), and mean normalized counts (baseMean). Genes listed meet a significance threshold of padj ≤ 0.05 and log2 fold change ≥ 0.5. Positive log2 fold change values indicate higher expression in target cell type group.

**Supplementary table 2: Differentially expressed genes (DEGs) for neuron cell types.** DEGs were calculated using a pseudobulk deseq2 approach that compared gene expression of each neuron cell type to all other neuron cell types (see Methods). The table reports adjusted p-values (padj) using BH correction for FDR, log2 fold change (log2FoldChange), standard error (lfcSE), raw p-values (pvalue), and mean normalized counts (baseMean). Genes listed meet a significance threshold of padj ≤ 0.05 and log2 fold change ≥ 0.5. Positive log2 fold change values indicate higher expression in target neuron cell type. Neurons for this group were defined as canonical neuron cell types of the CNS.

**Supplementary table 3: Differentially expressed genes (DEGs) for neuron developmental stages.** DEGs were calculated using a pseudobulk deseq2 approach that compared gene expression of each developmental stage (neuroblast, immature, and mature) to all other developmental stages (see Methods). The table reports adjusted p-values (padj) using BH correction for FDR, log2 fold change (log2FoldChange), standard error (lfcSE), raw p-values (pvalue), and mean normalized counts (baseMean). Genes listed meet a significance threshold of padj ≤ 0.05 and log2 fold change ≥ 0.5. Positive log2 fold change values indicate higher expression in target developmental stage.

**Supplementary table 4: Differentially expressed genes (DEGs) for neuron brain divisions.** DEGs were calculated using a pseudobulk deseq2 approach that compared gene expression of each brain division (telencephalon, diencephalon, mesencephalon, rhombencephalon) to all other brain divisions (see Methods). The table reports adjusted p-values (padj) using BH correction for FDR, log2 fold change (log2FoldChange), standard error (lfcSE), raw p-values (pvalue), and mean normalized counts (baseMean). Genes listed meet a significance threshold of padj ≤ 0.05 and log2 fold change ≥ 0.5. Positive log2 fold change values indicate higher expression in target brain division.

**Supplementary table 5: Differentially expressed genes (DEGs) for neuron brain regions.** DEGs were calculated using a pseudobulk deseq2 approach that compared gene expression of each brain region to all other brain regions (see Methods). The table reports adjusted p-values (padj) using BH correction for FDR, log2 fold change (log2FoldChange), standard error (lfcSE), raw p-values (pvalue), and mean normalized counts (baseMean). Genes listed meet a significance threshold of padj ≤ 0.05 and log2 fold change ≥ 0.5. Positive log2 fold change values indicate higher expression in target brain region.

**Supplementary table 6:** Differentially expressed genes (DEGs) between *A. mexicanus* populations within optic tectum (OT) neuron cell types. DEGs were calculated using a pseudobulk deseq2 approach within each OT neuron cell type (excitatory e2 or inhibitory i5), comparing cavefish to surface fish (see Methods). The table reports mean normalized counts (baseMean), log2 fold change (log2FoldChange), standard error (lfcSE), Wald statistic (stat), raw p-values (pvalue), adjusted p-values (padj; Benjamini–Hochberg corrected), and shrinkage-estimated log2 fold change (log2FoldChange_shrunk). Genes listed meet a significance threshold of padj ≤ 0.05. The table also reports whether the gene is enriched in cavefish or surface fish (significance), which reflects a positive log2 fold change for cavefish enrichment, with surface fish as the reference population. Additional table columns reflect entrez ID input (GeneID) into DAVID bioinformatics Gene ID conversion for *Astyanax mexicanus* (Species) to add full gene name (Gene.Name) and DAVID gene annotation (final_gene_symbol).

**Supplementary table 7: Differentially expressed genes (DEGs) for reclustered optic tectum (OT) cell types.** DEGs were calculated using a pseudobulk deseq2 approach that compared gene expression of each OT cell type to all other OT cell types (see Methods). The table reports adjusted p-values (padj) using BH correction for FDR, log2 fold change (log2FoldChange), standard error (lfcSE), raw p-values (pvalue), and mean normalized counts (baseMean). Genes listed meet a significance threshold of padj ≤ 0.05 and log2 fold change ≥ 0.5. Positive log2 fold change values indicate higher expression in target OT cell type.

**Supplementary table 8: Reclustered optic tectum (OT) wiring gene sets used for module-score analysis.** Gene sets were manually curated into four neuronal developmental wiring programs: patterning, route choice, target refinement, and synapse organization. For each wiring program, module scores were calculated for each OT reclustered cell type, then median module scores were z-scored and plotted as a heatmap for visualization (see SF5).

**Supplementary table 9: Differentially expressed genes (DEGs) between *A. mexicanus* populations within pineal neurons.** DEGs were calculated using a pseudobulk deseq2 approach within the pineal neuron cell type (e9), comparing cavefish to surface fish (see Methods). The table reports mean normalized counts (baseMean), log2 fold change (log2FoldChange), standard error (lfcSE), Wald statistic (stat), raw p-values (pvalue), adjusted p-values (padj; Benjamini–Hochberg corrected), and shrinkage-estimated log2 fold change (log2FoldChange_shrunk). Genes listed meet a significance threshold of padj ≤ 0.05. The table also reports whether the gene is enriched in cavefish or surface fish (significance), which reflects a positive log2 fold change for cavefish enrichment, with surface fish as the reference population. Additional table columns reflect entrez ID input (GeneID) into DAVID bioinformatics Gene ID conversion for *Astyanax mexicanus* (Species) to add full gene name (Gene.Name) and DAVID gene annotation (final_gene_symbol).

**Supplementary table 10: KEGG pathway enrichment results for pineal neuron differentially expressed genes (DEGs).** KEGG pathway enrichment analysis was performed on pineal neuron DEGs (ST9) between cave and surface populations. The table reports pathway category, KEGG term and ID, gene counts, enrichment statistics (PValue, Bonferroni, Benjamini, and FDR-adjusted p-values), fold enrichment, and associated gene lists. “Up in Cave” and “Up in Surface” indicate the direction of differential expression for genes contributing to each enriched pathway.

**Supplementary table 11: Differentially expressed genes (DEGs) for other, non-CNS neurons or glia cell types.** DEGs were calculated using a pseudobulk deseq2 approach that compared gene expression of each cell type to all other cell types (see Methods). The table reports adjusted p-values (padj) using BH correction for FDR, log2 fold change (log2FoldChange), standard error (lfcSE), raw p-values (pvalue), and mean normalized counts (baseMean). Genes listed meet a significance threshold of padj ≤ 0.05 and log2 fold change ≥ 0.5. Positive log2 fold change values indicate higher expression in target cell type.

**Supplementary table 12: Differentially expressed genes (DEGs) between *A. mexicanus* populations within pineal photoreceptors.** DEGs were calculated using a pseudobulk deseq2 approach within the pineal photoreceptor cell type (b7), comparing cavefish to surface fish (see Methods). The table reports mean normalized counts (baseMean), log2 fold change (log2FoldChange), standard error (lfcSE), Wald statistic (stat), raw p-values (pvalue), adjusted p-values (padj; Benjamini–Hochberg corrected), and shrinkage-estimated log2 fold change (log2FoldChange_shrunk). Genes listed meet a significance threshold of padj ≤ 0.05. The table also reports whether the gene is enriched in cavefish or surface fish (significance), which reflects a positive log2 fold change for cavefish enrichment, with surface fish as the reference population. Additional table columns reflect entrez ID input (GeneID) into DAVID bioinformatics Gene ID conversion for *Astyanax mexicanus* (Species) to add full gene name (Gene.Name) and DAVID gene annotation (final_gene_symbol).

**Supplementary table 13: Differentially expressed genes (DEGs) for glia cell types.** DEGs were calculated using a pseudobulk deseq2 approach that compared gene expression of each glial cell type to all other glial cell types (see Methods). The table reports adjusted p-values (padj) using BH correction for FDR, log2 fold change (log2FoldChange), standard error (lfcSE), raw p-values (pvalue), and mean normalized counts (baseMean). Genes listed meet a significance threshold of padj ≤ 0.05 and log2 fold change ≥ 0.5. Positive log2 fold change values indicate higher expression in target glial cell type.

**Supplementary table 14: Differentially expressed genes (DEGs) for radial glia subtypes.** DEGs were calculated using a pseudobulk deseq2 approach that compared gene expression of each radial glial subtype to all other subtypes (see Methods). The table reports adjusted p-values (padj) using BH correction for FDR, log2 fold change (log2FoldChange), standard error (lfcSE), raw p-values (pvalue), and mean normalized counts (baseMean). Genes listed meet a significance threshold of padj ≤ 0.05 and log2 fold change ≥ 0.5. Positive log2 fold change values indicate higher expression in target radial glial subtype.

**Supplementary table 15: Differentially expressed genes (DEGs) between *A. mexicanus* populations within microglia.** DEGs were calculated using a pseudobulk deseq2 approach within microglia, comparing cavefish to surface fish (see Methods). The table reports mean normalized counts (baseMean), log2 fold change (log2FoldChange), standard error (lfcSE), Wald statistic (stat), raw p-values (pvalue), adjusted p-values (padj; Benjamini–Hochberg corrected), and shrinkage-estimated log2 fold change (log2FoldChange_shrunk). Genes listed meet a significance threshold of padj ≤ 0.05. The table also reports whether the gene is enriched in cavefish or surface fish (significance), which reflects a positive log2 fold change for cavefish enrichment, with surface fish as the reference population. Additional table columns reflect entrez ID input (GeneID) into DAVID bioinformatics Gene ID conversion for *Astyanax mexicanus* (Species) to add full gene name (Gene.Name) and DAVID gene annotation (final_gene_symbol).

**Supplementary table 16: Differentially expressed genes (DEGs) between *A. mexicanus* populations within microglia calculated using cell-level analysis.** DEGs were calculated using FindMarkers in Seurat with the Model-based Analysis of Single-cell Transcriptomics (MAST) framwork, comparing cavefish to surface fish (see Methods).

The table reports raw p-values (p_val), average log2 fold change (avg_log2FC), the fraction of cells expressing each gene in cave and surface populations (pct.1 and pct.2, respectively), and adjusted p-values (p_val_adj). Genes listed meet a significance threshold of p_val_adj ≤ 0.05. Additional table columns reflect entrez ID input (GeneID) into DAVID bioinformatics Gene ID conversion for *Astyanax mexicanus* (Species) to add full gene name (Gene.Name) and DAVID gene annotation (final_gene_symbol).

## Notes

### Competing Interest Statement

The authors have declared no competing interest.

